# Conservation of the genetic interaction network between two yeast species

**DOI:** 10.1101/2025.09.21.677604

**Authors:** Carles Pons

**Affiliations:** Institute for Research in Biomedicine (IRB Barcelona), The Barcelona Institute for Science and Technology (BIST), 08028 Barcelona, Catalonia, Spain

## Abstract

Genetic interactions are essential to decipher the complex relationship between genotype and phenotype, which is key to addressing fundamental biology questions and developing therapies for human diseases. However, the scarcity of genetic interaction data in most species hinders research progress. Understanding the conservation of genetic interactions across species can partially bridge the current data gap. Previous conservation studies focused on subsets of genes and relied on the incomplete genetic networks available at the time, limiting the generalization of their findings. This study systematically quantifies the conservation of the genetic interaction networks between *Saccharomyces cerevisiae* and *Schizosaccharomyces pombe* using a comprehensive global genetic interaction map. Conservation is evaluated at three levels of increasing complexity: gene connectivity, individual genetic interactions, and genetic interaction profiles. Overall, the analyses show significant conservation across species at all levels, particularly among functionally related genes, 1:1 orthologs, and less diverged orthologs. However, the functional redundancy caused by gene duplication decreases cross-species conservation, resulting in lower connectivity, loss of individual interactions, and enrichment for trigenic interactions. This disruptive effect can be partially reversed upon divergence of the duplicated pair. Additionally, this work provides predictions for genetic interactions in *S. pombe* and trigenic interactions in *S. cerevisiae*. Together, the findings presented here offer a framework to explore the genetic interaction landscape in species lacking such experimental data, including cancer cell lines once large-scale genetic interaction data for human becomes available.

## INTRODUCTION

Phenotypes are rarely determined by single genetic variants and often arise from multiple contributing genetic factors (Hartman et al, 2001), contributing to the complexity of the genotype-to-phenotype problem (Dowell et al, 2010). Genetic interactions identify combinations of mutations that result in unexpected phenotypes (Costanzo et al, 2019), and can shed light on the intricate relationship between genetic variants and observable traits (Manolio et al, 2009; Zuk et al, 2012). In negative genetic interactions, the phenotype of the combined mutants is more deleterious than expected given the individual mutant phenotypes. Conversely, in positive genetic interactions, the combined phenotype is healthier than expected (Mani et al, 2008).

High-throughput genetic interaction screens were first pioneered in *Saccharomyces cerevisiae* (Tong et al, 2001, 2004), in which most gene pairs have already been screened to define the first global genetic interaction network in any species (Costanzo et al, 2016). This comprehensive map enabled accurate prediction of gene function, and outlined the functional architecture of the cell. Functionally related genes exhibited similar patterns of genetic interactions, with close functional relationships such as cocomplex membership showing higher similarity than distant relationships such as colocalization (Bandyopadhyay et al, 2008; Costanzo et al, 2010, 2016).

Genetic interactions have been studied in other model organisms but at a much smaller scale (Lehner et al, 2006; Dixon et al, 2008; Frost et al, 2012; Ryan et al, 2012; Fischer et al, 2015; Horlbeck et al, 2018; Heigwer et al, 2023; Billmann et al, 2025). Still, the general principles of genetic interaction networks seem similar across species. For instance, gene pairs within the same biological process are more likely to interact than unrelated pairs, and genes with stronger fitness defects tend to be more connected than genes without a phenotype (Byrne et al, 2007; Koch et al, 2012; Costanzo et al, 2016; Billmann et al, 2025). However, the limited amount of genetic interaction data outside *S. cerevisiae* hinders research progress in other species. This is particularly relevant in the study of human disease, since genetic interactions could reveal disease mechanisms (Wang et al, 2017; Fang et al, 2019), pinpoint specific weaknesses of cancer cells (Hartwell et al, 1997), and facilitate progress in personalized medicine (Hartman et al, 2001; Tutuncuoglu & Krogan, 2019).

Despite recent progress (Shen et al, 2017; Heigwer et al, 2023; Billmann et al, 2025), implementing large-scale genetic interaction screens in other species remains technically challenging. Additionally, the digenic space in some species is vast (in human it is ∼10 times larger than in *S. cerevisiae*), making efficient protocols crucial. Thus, in most species the exhaustive screening of genetic interactions is not likely in the near future. An alternative to experimental screens entails using the global genetic interaction network from *S. cerevisiae* to infer interactions in other species via orthology relationships. Previous studies found partial conservation across species for subsets of genes, like those in particular pathways or protein complexes (Dixon et al, 2008; Roguev et al, 2008; Ryan et al, 2012). However, these analyses considered the partial genetic interaction networks available at the time, and mostly focused on 1:1 orthologs, neglecting duplicated orthologs. These constraints limit the generalization of the findings.

This study systematically characterizes the conservation between the genetic interaction networks in *S. cerevisiae* and *Schizosaccharomyces pombe* using the datasets with the largest coverage in any species, including a comprehensive global map. Conservation is evaluated at three levels of increasing complexity: gene connectivity, individual genetic interactions, and genetic interaction profiles. Overall, the presented analyses quantify the conservation rates across species, identify the conditions in which conservation is most likely, and provide predictions which can be readily tested.

## RESULTS AND DISCUSSION

Genetic interaction data for *S. cerevisiae* was generated from ∼23 million double mutants including ∼90% of the ∼6,000 genes in that species (Costanzo et al, 2016). In total, ∼550,000 negative and ∼350,000 positive genetic interactions were identified. The *S. pombe* data contained measurements for ∼1.6 million double mutants including ∼50% of the ∼5,000 genes in that species, resulting in ∼25,000 negative and ∼12,000 positive genetic interactions (Ryan et al, 2012).

### Conservation of genetic interaction density

The frequency of genetic interactions of a given gene, known as genetic interaction density, is calculated by dividing its number of genetic interactions by the number of screened gene pairs. This metric summarizes the network connectivity of a gene and reflects its importance in the cell. Indeed, genes with strong fitness defects tend to have a higher negative genetic interaction density than genes with weak phenotypes (Koch et al, 2012).

Here, to evaluate if gene connectivity in the genetic interaction network was conserved across species, the genetic interaction densities in *S. cerevisiae* and *S. pombe* were compared. The correlation of densities was moderate (**Figure 1A**), and slightly higher for the negative than for the positive genetic interaction network (0.22 and 0.19, respectively). Correlation was stronger between 1:1 orthologs, reaching 0.40 in the negative genetic interaction network, in line with a previous study (Koch et al, 2012), and weaker between duplicated genes (**Figure 1B**).

**Figure 1.**
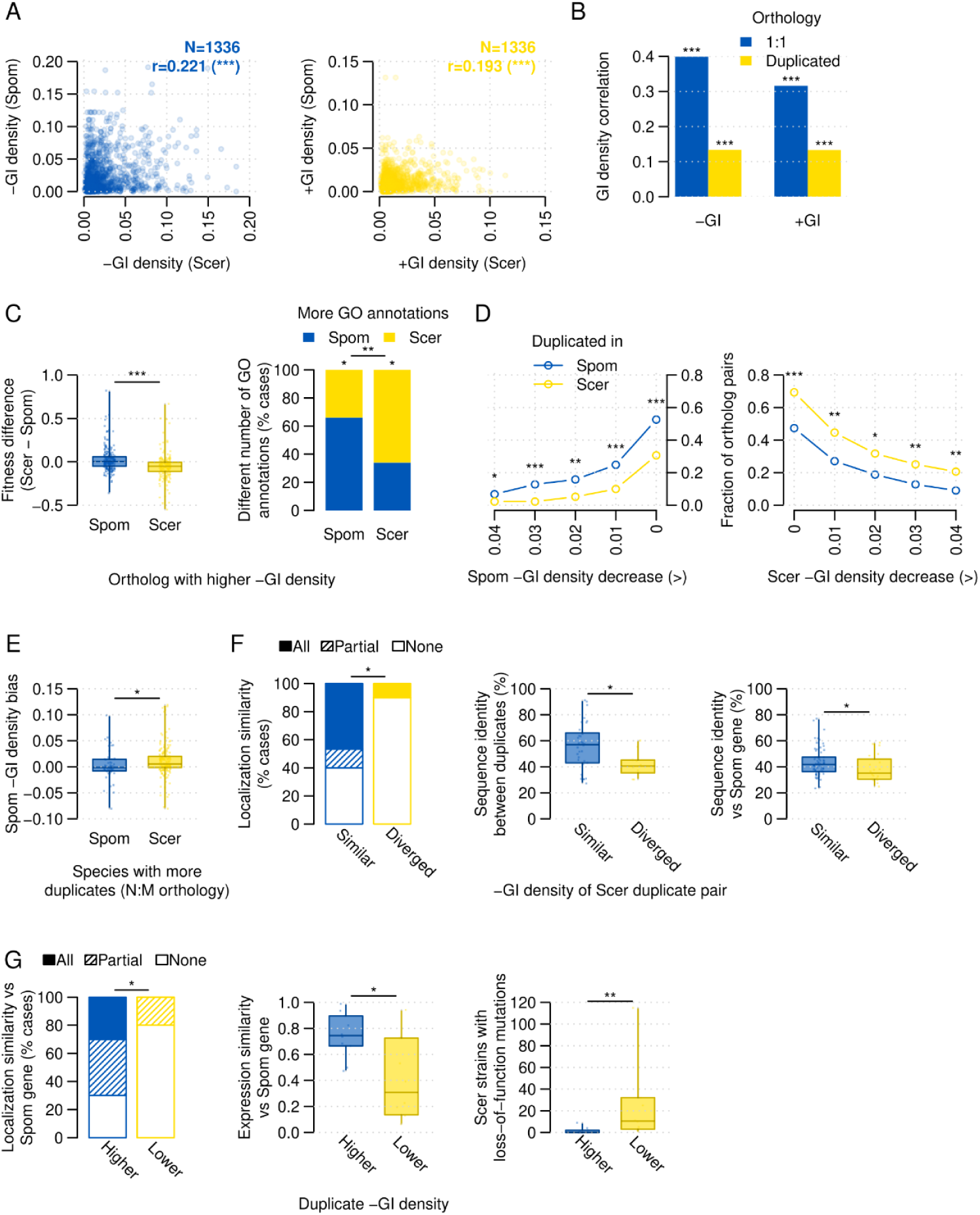
Conservation of the connectivity in the genetic interaction network. A: scatter plot of negative (left) and positive (right) genetic interaction densities in S. cerevisiae and S. pombe. B: barplot showing the correlation of the genetic interaction densities in S. cerevisiae and S. pombe for 1:1 orthologs and for duplicated genes. C: (left) boxplot showing the difference in single mutant fitness between S. cerevisiae genes and the S. pombe orthologs; (right) stacked barplot showing the fraction of orthologs with more GO annotations in S. pombe than in S. cerevisiae, and vice versa. Ortholog pairs are split by the species in which they have a higher negative genetic interaction density. D: plot showing the fraction of ortholog pairs with lower negative genetic interaction density than expected in (left) S. pombe and (right) S. cerevisiae at different cutoffs. Only pairs involving duplicated pairs in either species are shown. E: boxplot showing the negative genetic interaction bias for ortholog pairs involving N:M orthology relationships. Negative and positive values identify cases with fewer and more interactions than expected in S. pombe, respectively. F: (left) plot showing the fraction of duplicate pairs in S. cerevisiae with complete, partial, or no similarity in cellular localization; (center) boxplot showing the sequence identity between duplicate pairs; (right) boxplot showing the interspecies sequence identity between the S. cerevisiae duplicates and the corresponding single ortholog in S. pombe. All plots compare duplicate pairs in S. cerevisiae in which: i) both genes have lower negative genetic interaction density than expected given the density of the S. pombe ortholog, and ii) the rest of duplicate pairs. G: (left) plot showing the fraction of duplicated genes in S. cerevisiae with complete, partial, or no similarity in cellular localization with the corresponding single ortholog in S. pombe; (center) boxplot showing the expression similarity of duplicated genes in S. cerevisiae with the corresponding single ortholog in S. pombe; (right) boxplot showing the number of S. cerevisiae strains in which duplicated genes in S. cerevisiae have a loss-of-function mutation. Only duplicate pairs with a divergent negative genetic interaction density are shown. Within each duplicate pair, genes are split by their negative genetic interaction density. Statistical significance was calculated by Fisher’s exact (D, C-right), binomial (C-right) and Mann-Whitney U tests (C-left, E, F, and G). ns: not significant; *: p < 0.05; **: p < 0.005; ***: p < 0.0005.

Orthologs with higher density in *S. pombe* than in *S. cerevisiae* exhibited stronger fitness defects in *S. pombe*. Conversely, genes more connected in *S. cerevisiae* tended to have stronger phenotypes in that species (**Figure 1C**). Thus, changes in negative genetic interaction density across species were in agreement with the severity of the individual phenotypes. Similarly, 1:1 ortholog pairs generally had more GO annotations (Gene Ontology Consortium et al, 2023) in the species with the highest density (**Figure 1C**).

To evaluate the effect of gene duplication across species, genes with a single copy in one of the species and duplicated in the other were selected. Ortholog pairs with a lower negative genetic interaction density in *S. cerevisiae* than expected (see Methods) were enriched for genes duplicated in that species. The same pattern was observed in genes with lower density in *S. pombe* (**Figure 1D**). Thus, upon duplication, genes exhibited fewer negative genetic interactions than expected given the connectivity of their single-copy ortholog gene, likely because of the functional redundancy of duplicates. The more complex N:M orthology relationships, in which multiple genes in one species map to multiple genes in the other, showed the same trend. In those cases, ortholog sets with more duplicates in *S. pombe* exhibited a lower negative genetic interaction density in that species than expected (**Figure 1E**). It is worth noting that some of the observed trends in connectivity changes across species are in line with previous reports focused on a single species, such that singletons, multifunctional genes, and genes with a strong phenotype were already reported to have more genetic interactions than duplicates, genes with a single function, and genes with a weak phenotype, respectively (VanderSluis et al, 2010; Koch et al, 2012; Costanzo et al, 2016; Kuzmin et al, 2020).

Next, gene duplicates in *S. cerevisiae* with a single ortholog in *S. pombe* were split by whether they had similar or diverged negative genetic interaction densities in *S. cerevisiae*. The duplicates with similar densities showed higher similarity in sequence and cellular localization pattern, and in sequence to the *S. pombe* single ortholog (**Figure 1F**). Thus, a higher rate in the acquisition of mutations or changes in localization patterns is reflected in a more dissimilar connectivity in the genetic interaction network. Additionally, in duplicate pairs that diverged in negative genetic interaction density, the gene with the higher connectivity tended to be more similar in localization and expression patterns to the *S. pombe* single ortholog, and to have fewer loss-of-function mutations across a panel of 1,011 S. cerevisiae strains (Peter et al, 2018) (**Figure 1G**).

### Conservation of genetic interactions

Considering the largest studies in both species (Ryan et al, 2012; Costanzo et al, 2016), out of the 12.5m gene pairs in *S. cerevisiae* for which genetic interactions have been screened, 36% correspond to genes with orthologs in *S. pombe*. However, most of those pairs have not been screened in that species, and only 13% gene pairs have genetic interaction data available. In total, ∼600k gene pairs with genetic interaction data in *S. cerevisiae* have also ortholog pairs screened for interactions in *S. pombe*, which enabled conservation analyses.

Here, the cross-species conservation of a given *S. cerevisiae* genetic interaction was evaluated by inspecting all possible ortholog pairs in *S. pombe* (see Methods; **Figure 2A**). For instance, if both *S. cerevisiae* genes had one ortholog in *S. pombe*, only one gene pair was examined in that species. However, if both genes had two orthologs, then four different pairs were examined (**Figure 2A**). Importantly, an interaction between any of the ortholog pairs was considered enough to indicate genetic interaction conservation. In total, 958 negative (5.5%) and 279 positive (2.9%) genetic interactions identified in *S. cerevisiae* were conserved in *S. pombe*, 1.9 and 1.6 times more than expected, respectively (p < 0.0005, **Figure 2B**). Conservation was higher when both genes had 1:1 orthologs (7.0% for negative and 3.2% for positive genetic interactions, 3.0 and 2.6 fold enrichment, respectively), and decreased substantially when both genes were duplicated in either species (**Figure 2C**). The result for 1:1 orthologs is consistent with the rates previously found in a smaller dataset (9% and 4.2%, respectively; see supplementary information in (Ryan et al, 2012)). Additionally, extreme interactions were more often conserved than weaker ones (**Figure 2D**).

**Figure 2.**
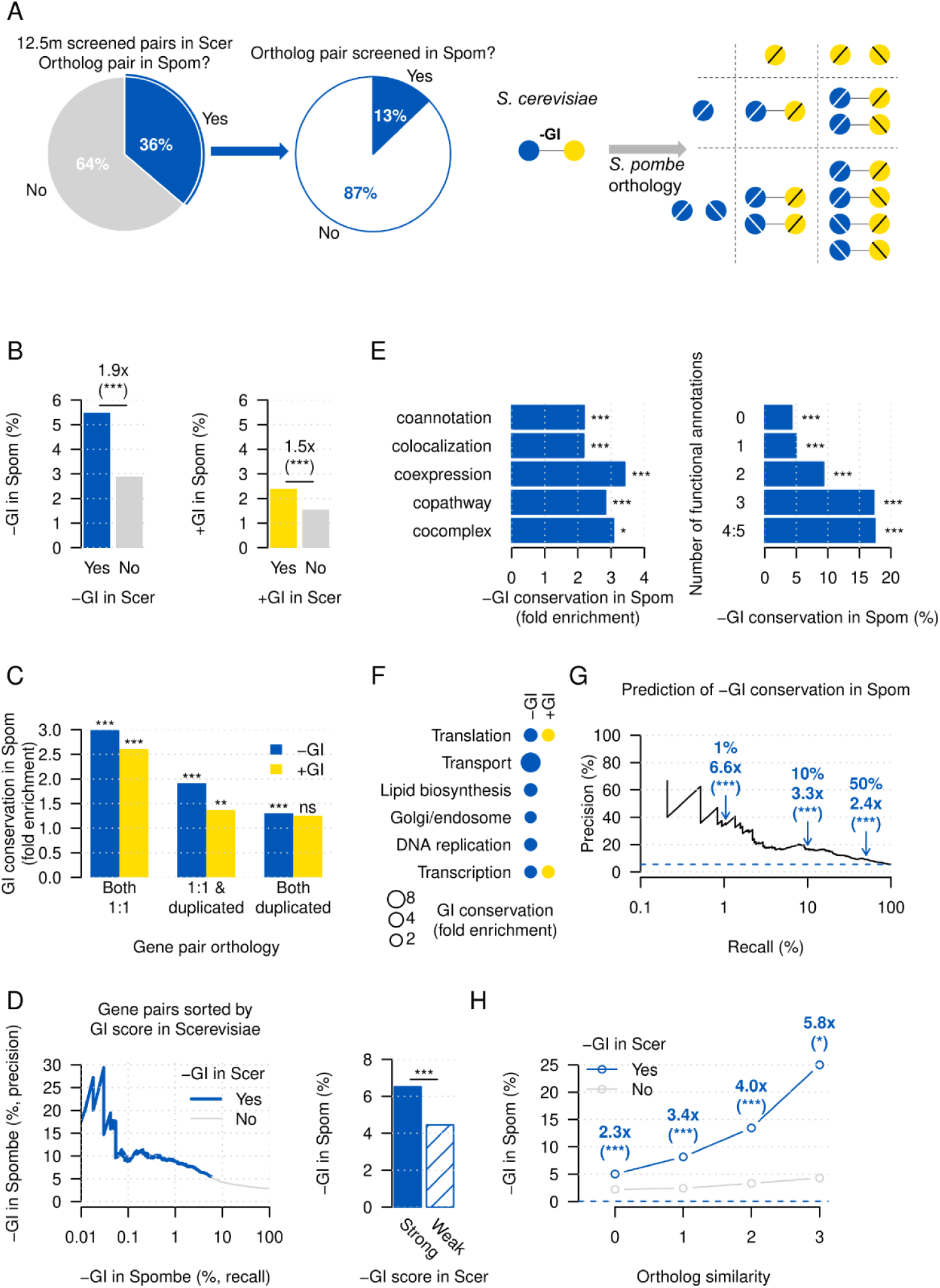
- Conservation of genetic interactions. A: (left) pie chart showing the fraction of gene pairs in S. cerevisiae with genetic interaction data that have orthologs in S. pombe; (center) pie chart showing the fraction of ortholog pairs in S. pombe for which genetic interaction data is available; (right) diagram showing how genetic interaction conservation is calculated between S. cerevisiae and S. pombe across different orthology relationships. For S. cerevisiae genes with a single ortholog in S. pombe, only one ortholog pair has to be evaluated in that species. For S. cerevisiae genes with two orthologs in S. pombe, four ortholog pairs have to be evaluated. B: barplot showing the fraction of interacting and non-interacting pairs in S. cerevisiae that have an interaction in S. pombe, for (left) negative and (right) positive genetic interactions. C: barplot showing the fold enrichment in the conservation of genetic interactions for gene pairs grouped by their orthology relationship between S. cerevisiae and S. pombe. D: (left) precision recall curve for the identification of conserved negative genetic interactions in S. pombe for gene pairs in S. cerevisiae sorted by their genetic interaction score; (right) barplot showing the fraction of conserved negative genetic interactions in S. pombe for negative genetic interactions in S. cerevisiae grouped by the interaction strength. E: barplots showing the fold enrichment in the conservation of negative genetic interactions in S. pombe for gene pairs in S. cerevisiae grouped by (left) different functional relationships, and (right) number of functional relationships. F: dotplot showing the fold enrichment in the conservation of genetic interactions for broad functional classes. Only classes with significant enrichments (p < 0.05) are shown. G: precision recall curve for the prediction of conserved negative genetic interactions in S. pombe using a random forest. Dots identify the performance at 1%, 10%, and 50% recall. H: plot showing the fraction of conserved negative genetic interactions in S. pombe for gene pairs in S. cerevisiae divided in four different levels of similarity to their corresponding 1:1 orthologs. Statistical significance was calculated by Fisher’s exact (B, C, D, F, G, H). ns: not significant; *: p < 0.05; **: p < 0.005; ***: p < 0.0005.

Next, gene pairs were annotated using five functional standards in *S. cerevisiae*. Interactions of functionally related pairs in *S. cerevisiae* were more frequently found in *S. pombe* (**Figure 2E**). For instance, interactions involving gene pairs coding for the same protein complex were 3.1 times more likely to be conserved than expected (p < 0.0005). Moreover, support by more functional relationships was associated with higher conservation rates, reaching 17.5% for interacting pairs annotated to four or more functional standards, compared to 4.3% for pairs absent in all the standards (**Figure 2E**). Finally, rates increased for genes involved in DNA replication, as in a recent study (Billmann et al, 2025), translation, transcription, and other functions (**Figure 2F**).

Using functional relationships, interaction strength, and orthology data of negative genetic interactions in *S. cerevisiae*, a random forest model was trained for the prediction of conserved interactions in *S. pombe*. The resulting predictions were enriched for negative genetic interactions in *S. pombe*, reaching conservation rates of 36% and 17% among the top 1% and top 10% ranked pairs, 6.6 and 3.3 times more than expected, respectively (p < 0.0005, **Figure 2G**), with an AUC of 0.63. These predictions for *S. pombe* are available in **Table S1** and can be used as a starting point for downstream analyses in that species.

To evaluate the relationship between ortholog divergence and interaction conservation, gene pairs in *S. cerevisiae* were classified by the similarity to their 1:1 orthologs in *S. pombe*. Three different metrics were used, including sequence identity, expression similarity, and cellular localization similarity (see Methods). Interactions of the most similar gene pairs across species were more often conserved, reaching 25% of the interactions, compared to only 5% when there was no gene similarity across species (p < 0.05, **Figure 2G**). This increasing trend in the conservation rate shows that less diverged gene pairs are more likely to maintain a similar function and gene relationships across species. This similarity may be lost because of changes in sequences, expression and localization patterns, which would result in a different pattern of genetic interactions between species. Addition of these divergence metrics to the conservation predictor improved the AUC to 0.66 for 1:1 orthologs.

### Conservation of negative genetic interactions when one gene duplicates

Upon gene duplication, the resulting paralogs functionally buffer each other, masking the original single mutant phenotype (Ohno, 1970; VanderSluis et al, 2010). Their phenotype is only revealed when both duplicates are simultaneously compromised. This buffering can be conserved across long evolutionary distances. For instance, phenotypes of single mutants in *S. cerevisiae* and *S. pombe* correlate for 1:1 orthologs, but not for duplicated genes (**Figure 3A**). In contrast, phenotypes of single mutants in *S. pombe* and the corresponding double mutants of duplicate pairs in *S. cerevisiae* still significantly correlate (**Figure 3A**). Thus, in a gene pair with a negative genetic interaction, a duplication in one gene would result in the loss of the interaction since the new paralogs functionally back up each other (**Figure 3B**). Also, upon duplication, the negative digenic interaction could become a trigenic one (Kuzmin et al, 2020) since the phenotype would only be revealed when compromising the three genes (**Figure 3B**). Divergence over time between the duplicates can compromise their functional redundancy and restore the digenic interaction for one of the duplicates.

**Figure 3.**
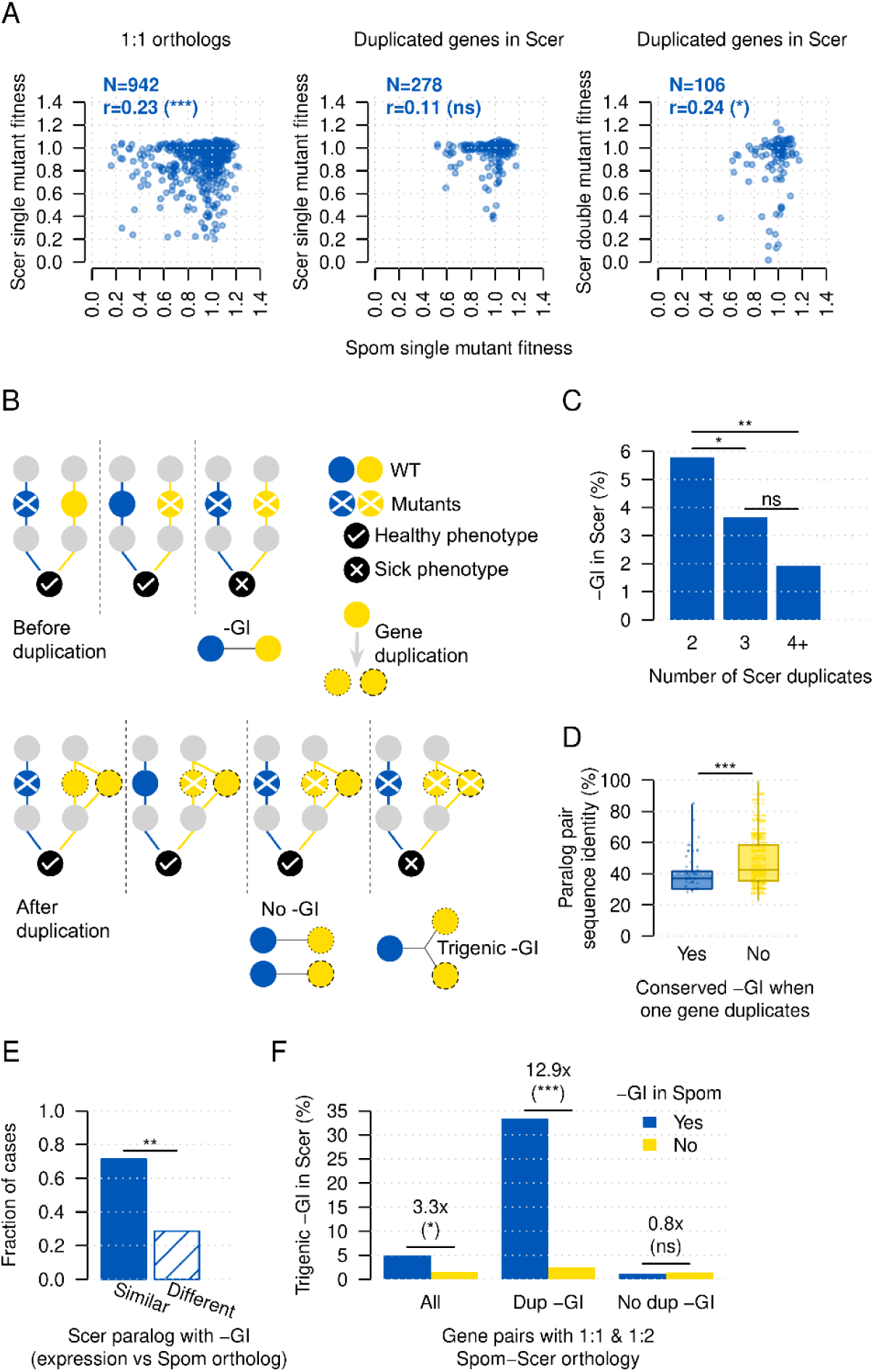
- Conservation of genetic interactions after duplication events. A: scatter plot of single mutant fitness in S. pombe and: (left) the single mutant fitness in S. cerevisiae for 1:1 orthologs; (center) the individual single mutant fitness in S. cerevisiae for each duplicated gene; (right) the double mutant fitness in S. cerevisiae of the duplicated pair. Only orthology relationships resulting in two duplicated genes in S. cerevisiae were considered. B: diagram showing how gene duplication can affect a negative digenic interaction and change into a negative trigenic interaction. C: barplot showing the fraction of gene pairs in S. cerevisiae that conserve the negative genetic interaction of the corresponding gene pair in S. pombe. Only S. pombe gene pairs with a negative genetic interaction in which one gene has a 1:1 ortholog and the other is duplicated in S. cerevisiae were selected. Gene pairs in S. cerevisiae are grouped by the total number of paralogs of the duplicated gene. D: boxplot showing the sequence identity between duplicated gene pairs in S. cerevisiae. Only S. pombe gene pairs with a negative genetic interaction in which one gene has a 1:1 ortholog and the other is duplicated in S. cerevisiae were selected. The interaction was considered to be conserved if any of the two duplicates and the corresponding 1:1 ortholog had a negative genetic interaction in S. cerevisiae. E: barplot showing the fraction of cases in which the S. cerevisiae paralog that has the conserved negative genetic interaction with the 1:1 ortholog has a more similar or divergent expression level compared to the corresponding S. pombe single ortholog. F: barplot showing the fraction of negative trigenic interactions for sets of three genes in S. cerevisiae in which one gene has a 1:1 ortholog in S. pombe and the other two are duplicates sharing a single ortholog in that species. “Dup -GI” and “No dup -GI” identify cases in which the duplicated genes in S. cerevisiae have and do not have a negative genetic interaction, respectively. Statistical significance was calculated by Fisher’s exact (C, F), Mann-Whitney U tests (D), and binomial (E). ns: not significant; *: p < 0.05; **: p < 0.005; ***: p < 0.0005.

Here, to better characterize the genetic interaction conservation dynamics among duplicates, *S. pombe* gene pairs with a negative interaction were selected, in which one gene had a 1:1 ortholog in *S. cerevisiae* and the other was duplicated. Then, *S. pombe* gene pairs were grouped by the number of duplicates in *S. cerevisiae*, and the corresponding *S. cerevisiae* gene pairs were evaluated for interaction conservation. *S. cerevisiae* gene pairs in which the duplicated gene had only one other paralog were more likely to conserve the interaction than if the duplicated gene had two or more paralogs (p < 0.05, **Figure 3C**).

Next, the subset of pairs in which the duplicated gene had only two orthologs in *S. cerevisiae* were selected, such that each *S. pombe* gene pair mapped to two *S. cerevisiae* gene pairs (totaling three different *S. cerevisiae* genes). In this set, a negative genetic interaction in *S. pombe* was more often conserved in any of the two *S. cerevisiae* gene pairs if the duplicates had diverged more in sequence (p < 0.0005, **Figure 3D**). This suggests that duplicate pairs with more similar sequences are more likely to be functionally redundant, therefore masking the interaction. However, as sequences and functions diverge between the duplicates, redundancy may be lost and any of the gene pairs would be more likely to restore the interaction. To determine whether the gene pair conserving the interaction could be predicted, the expression levels of the orthologs were compared. Interestingly, in 71% of the cases, the *S. cerevisiae* duplicate with an expression level more similar to the *S. pombe* ortholog was the one conserving the interaction (p < 0.005, **Figure 3E**).

To test if a negative digenic interaction in *S. pombe* could become trigenic in *S. cerevisiae* upon the duplication of one of the genes, data from recent large-scale trigenic studies was queried (Kuzmin et al, 2018, 2020). The corresponding sets of three *S. cerevisiae* genes were enriched for trigenic interactions (3.3 fold enrichment, p < 0.05, **Figure 3F**). Notably, this signal was dominated by cases in which the *S. cerevisiae* duplicates also exhibited a negative genetic interaction. In these cases, the rate of negative trigenic interactions was 13 times higher than expected (p < 0.0005), in agreement with a study that reported negatively interacting paralog pairs having more negative trigenic interactions than the remaining paralogs (Kuzmin et al, 2020). Cases in which the *S. cerevisiae* duplicates did not exhibit a negative digenic interaction were not enriched for trigenic interactions (**Figure 3F**). Thus, a combination of negative genetic interaction data in *S. pombe* and *S. cerevisiae*, and orthology relationships, can be used for the prediction of negative trigenic interactions in *S. cerevisiae*. **Table S2** contains such predictions for 176 sets of genes.

### Conservation of genetic interaction profile similarities

The set of genetic interactions of a gene is called genetic interaction profile (Tong et al, 2004). These profiles contain information on the cellular response to mutants in a specific genetic background, and are rich and informative phenotypic fingerprints (Bandyopadhyay et al, 2008; Costanzo et al, 2010). Genes with similar genetic interaction profiles identify strong functional relationships and have been previously used for the functional characterization of genes (Costanzo et al, 2016).

Here, genetic interaction profile similarities in *S. cerevisiae* and *S. pombe* were compared to evaluate if the profile similarity network was conserved across species. The global correlation was very poor (r=0.03, **Figure 4A**), likely because most gene pairs have non-significant similarities ∼0 which are not informative. Indeed, when considering only pairs with high similarity (top 10%, 1%, and 0.1% values), there was a strong agreement between both species (1.2, 7.2, and 91.7 fold enrichment, p < 0.0005, **Figure 4B**). As an example, for each gene, the most similar interaction profile in each network was selected. In 6.6% of the cases, the same gene was the most similar in both species (41.2 fold enrichment, p < 0.0005, **Figure 4C**). Additionally, in 15.4% and 25.0% of cases, the most similar gene in *S. cerevisiae* was among the top 1% and top 10% similarities in *S. pombe* (14.6 and 2.5 fold enrichment, respectively, p < 0.0005, **Figure 4C**). Gene duplication strongly affected the conservation of high profile similarities. Among the gene pairs with a high similarity in both species, the presence of one or two duplicated genes was 4.5 and 9.3 times lower than expected (p < 0.0005, **Figure 4D**). Conversely, pairs with 1:1 orthologs were 2.2 times more prevalent than expected.

**Figure 4.**
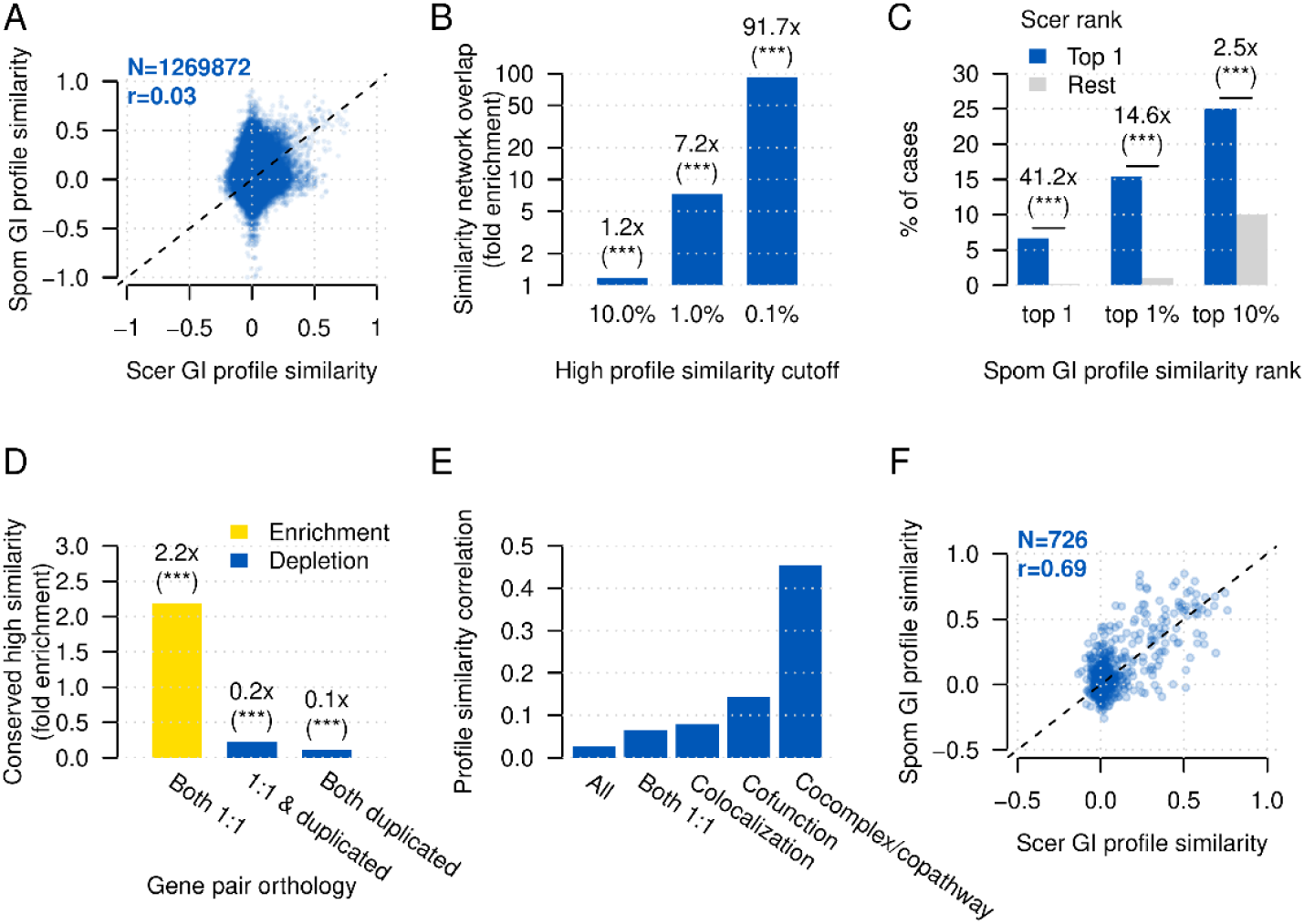
- Conservation of the genetic interaction profile similarity network. A: scatter plot of the genetic interaction profile similarity in S. cerevisiae and in S. pombe. B: barplot showing the fold enrichment in the overlap of the genetic interaction profile similarity networks in S. cerevisiae and S. pombe after applying different similarity cutoffs. C: barplot showing the fraction of 1:1 orthologs where their most similar genetic interaction profile in S. cerevisiae and S. pombe corresponds to the same gene. Also, cases where the most similar profile in S. cerevisiae rank among the top 1% and top 10% similarities in S. pombe are shown. D: barplot showing the fold enrichment for the conservation of high genetic interaction profile similarity between S. pombe and S. cerevisiae for gene pairs in which: i) both genes have a 1:1 ortholog; ii) one gene has a 1:1 ortholog and the other one is duplicated in S. cerevisiae; iii) both genes are duplicated in S. cerevisiae. E: barplot showing the correlation between the genetic interaction profile similarities in S. cerevisiae and S. pombe for all gene pairs, and gene pairs: with 1:1 orthologs, localized in the same cellular compartment in S. cerevisiae, with the same broad function in S. cerevisiae, and in the same molecular pathway or protein complex in S. cerevisiae. F: scatter plot showing the correlation between the genetic interaction profile similarities in S. cerevisiae and S. pombe for the subset of gene pairs with 1:1 orthologs and in the same molecular pathway or protein complex. Statistical significance was calculated by Fisher’s exact (B, C, D). ns: not significant; *: p < 0.05; **: p < 0.005; ***: p < 0.0005.

The correlation of genetic interaction profile similarities across species was higher for some sets of gene pairs, like those with 1:1 orthologs (r=0.07). Also, correlation was stronger for functionally related genes, with higher values for members of the same molecular pathway or protein complex (r=0.46), and lower values for less specific relationships like colocalization (r=0.08, **Figure 4E**). Notably, the set of 726 copathway and cocomplex gene pairs in *S. cerevisiae* with 1:1 orthologs in *S. pombe* had a particularly high agreement, reaching r=0.69 and highlighting that genetic interaction profile similarities are strongly conserved for specific gene sets (**Figure 4F**).

### Comparison of genetic interaction profiles across species

Ortholog pairs with conserved biological roles tend to exhibit similar patterns of molecular and functional interactions across species (Yu et al, 2004). Since genetic interaction profiles represent phenotypic signatures, similar profiles across species may identify cases in which assays in one species could be informative in another. However, the different experimental setups and scoring procedures used to derive genetic interaction data in *S. cerevisiae* and *S. pombe*, together with incomplete screening spaces, could complicate such comparisons.

Here, to determine if similar interaction profiles across species could reflect functional conservation, the interspecies genetic interaction profile similarities between all *S. cerevisiae* and all *S. pombe* genes were computed (see Methods). Reassuringly, ortholog pairs showed higher correlations than unrelated pairs (p < 0.0005, **Figure 5A**), particularly at high similarity cutoffs. For instance, orthologs were 9.3 times more likely to have a similarity above 0.3 than other gene pairs (p < 0.0005, **Figure 5A**). Next, orthologs with high interspecies similarity were selected and compared to the remaining orthologs. Characterization using Qarles (Pons, 2025) showed that they tended to code for protein complexes (Meldal et al, 2021), to exhibit a stronger fitness defect (Costanzo et al, 2016), and to have a lower mRNA halflife (Chan et al, 2018) in *S. cerevisiae* (p < 0.05, **Figure 5B**). Also, these genes were more often involved in transcription and chromosome segregation (p < 0.05, **Figure 5C**), and had more similar cellular localization patterns across species (p < 0.05, **Figure 5B**).

**Figure 5.**
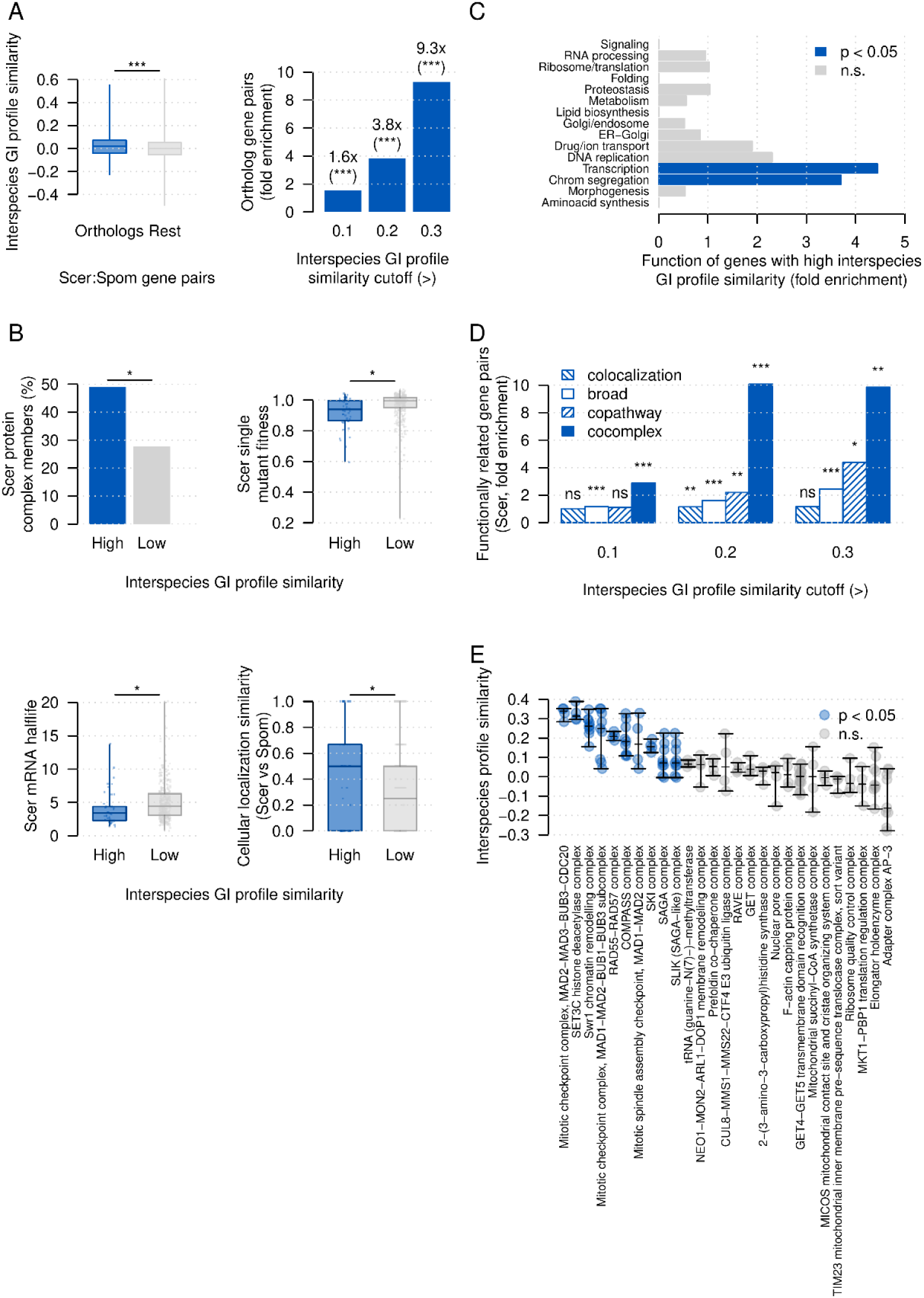
Interspecies genetic interaction profile similarity. A: (left) boxplot showing the interspecies genetic interaction profile similarity for ortholog and non-ortholog gene pairs; (right) barplot showing the fold enrichment for ortholog gene pairs among profile similarities above three different cutoffs. B: comparison between 1:1 ortholog gene pairs with high and low interspecies genetic interaction profile similarity: (top left) barplot showing the fraction of protein complex members in S. cerevisiae; (top right) boxplot showing the single mutant fitness in S. cerevisiae; (bottom left) boxplot showing mRNA halflife in S. cerevisiae; (bottom right) boxplot showing the Jaccard index of the interspecies cellular localization similarity. C: barplot showing the fold enrichment for S. cerevisiae broad functional annotations among ortholog gene pairs with high interspecies similarity. D: barplot showing the fold enrichment for functionally related gene pairs with high interspecies genetic interaction profile similarity at different stringency cutoffs. Functional data from S. cerevisiae. E: plot showing the interspecies similarity for S. cerevisiae protein complexes with data available for at least two subunits. Black lines show the minimum, maximum, and median value per complex. Statistical significance was calculated by Fisher’s exact (A right, B top left, C, D), and Mann-Whitney U tests (A left, B, E). ns: not significant; *: p < 0.05; **: p < 0.005; ***: p < 0.0005.

Besides orthologs, gene pairs with a functional relationship were enriched for high interspecies profile similarities, particularly members of the same protein complex (p < 0.05, **Figure 5D**).

Specific complexes exhibited higher similarities, suggesting that their biological role could be highly conserved (**Figure 5E**), including the Mitotic checkpoint complex, the Swr1 chromatin remodelling complex, the COMPASS complex, and the SAGA complex.

## CONCLUDING REMARKS

Beyond *S. cerevisiae*, genetic interaction data is scarce and, except for a handful of model organisms (Billmann et al, 2025), will continue to be in the near future. An alternative to the experimental screening of genetic interactions is to use a reference network and infer interactions in other species by orthology relationships. Previous conservation analyses of genetic interactions focused on partial networks, small sets of genes, and 1:1 orthologs (Dixon et al, 2008; Roguev et al, 2008; Ryan et al, 2012), limiting the scope of the findings. The work presented here extends previous studies by: i) leveraging a global genetic interaction map in *S. cerevisiae*; ii) evaluating conservation at three levels of increasing complexity; iii) focusing on the effects of gene duplication on conservation.

Across all analyses, gene pairs with 1:1 orthologs consistently exhibited higher conservation rates than gene pairs with other orthology relationships, as expected for genes more likely to have similar functions. Conservation was also higher for functionally related gene pairs, and less diverged ortholog pairs. Importantly, ortholog divergence was not only determined by sequence but also by changes in cellular localization and PPI partners, reflecting the complexity of functional variation. Additionally, the observed changes in connectivity between species were in line with previously reported trends within a single species, highlighting the general design principles of genetic interaction networks (Koch et al, 2012; Ryan et al, 2012; Costanzo et al, 2016).

The disruptive effect of gene duplication in genetic networks was significant in all cross-species comparisons. Functional redundancy between duplicates was linked to a decrease in gene connectivity, and the loss of individual genetic interactions, consistent with findings from previous single-species analyses (VanderSluis et al, 2010; Kuzmin et al, 2020). Duplicate pairs with asymmetric connectivity were more divergent in sequence and cellular localization than duplicates with similar connectivity. The more connected gene duplicate was more similar in sequence and localization pattern to the corresponding single ortholog than the other duplicate. Additionally, negative genetic interactions in *S. pombe* in which one gene was duplicated in *S. cerevisiae* were enriched for trigenic interactions in that species. Thus, the functional buffering caused by duplication events enabled the prediction of higher-order genetic interactions.

The moderate conservation rates reported here represent a lower estimate given the modest reproducibility of genetic interaction screens and the different methods (experimental and computational) used to derive the analyzed genetic interaction data. Additionally, the ∼400 million years of divergence between *S. cerevisiae* and *S. pombe* expectedly result in extensive functional rewiring, particularly for duplicated genes. Thus, higher conservation is expected between more closely related species and among cancer cell lines. Importantly, the rewiring of the genetic interaction network can be partially predicted by quantifying divergence between orthologs and duplicates.

Comprehensive genetic interaction data may soon be available for human cell lines and particular species. However, complete maps for every human individual, cancer type, or species are unlikely in the near future. Thus, determining how reference maps can translate to particular genetic variants is critical. Such knowledge would boost the implementation of personalized medicine, enable the identification of specific cancer vulnerabilities, and increase our understanding of the fundamental biology in other organisms. This study aims to modestly contribute to that goal by comprehensively characterizing the conservation of the genetic interaction networks between two yeast species. The use of a global reference genetic network makes the findings generalizable, providing a framework which can be applied to different species.

## MATERIALS & METHODS

### Correlation of genetic interaction densities (1A, 1B)

Negative genetic interaction density was calculated for each gene by dividing its number of negative genetic interactions by the number of screened pairs. For *S. cerevisiae*, density was calculated for all array genes after applying the intermediate cutoff. For gene pairs screened more than once, the pair was considered to interact if 50% or more of the tested combinations resulted in a negative genetic interaction. For *S. pombe*, interaction density was calculated for all array genes. DAmP alleles were not considered in this study for any of the datasets. The same procedure was used to calculate positive genetic interaction densities. Next, a table was generated including all orthology relationships, such that 1:1 orthologs contributed one gene pair, N:1 and 1:N orthologs contributed N pairs, and N:M orthologs contributed NxM pairs. Only gene pairs with genetic interaction data available in both species were kept. The Spearman’s correlation coefficient was calculated for: i) all pairs; ii) pairs in which both genes had 1:1 orthologs; iii) pairs in which at least one of the genes was duplicated in either species. Orthology mappings were retrieved using Panther 16.1 (Mi et al, 2021).

### Deviation from the expected interaction density (1C, 1D, 1E)

For each ortholog gene pair, the interspecies genetic interaction density ratio was calculated by dividing the interaction density in *S. pombe* by the density in *S. cerevisiae*. The median ratio among 1:1 ortholog pairs was used as the expected density relationship between both species. Consequently, the expected density in *S. pombe* was calculated by multiplying that median ratio by the interaction density in *S. cerevisiae*. Then, the density deviation was calculated by subtracting the expected density from the real density in *S. pombe*, in which negative values identified cases with a lower *S. pombe* density than expected, and positive values identified cases with higher density in *S. pombe* than expected.

The interspecies fitness difference was calculated by subtracting the single mutant fitness in *S. pombe* (Koch et al, 2012) from the fitness in *S. cerevisiae* (Costanzo et al, 2016), such that negative values identified cases with a lower fitness (i.e. stronger fitness defect) in *S. cerevisiae*. Fitness differences between 1:1 orthologs were grouped by the deviation from the expected interaction density in *S. pombe*: ortholog pairs in which the observed *S. pombe* interaction density was (i) higher or (ii) lower (i.e. higher *S. cerevisiae* density) than the expected density. The interspecies fitness difference of the two sets were compared and the statistical significance was calculated by Mann-Whitney U tests. For the same two sets of 1:1 orthologs, the number of GO SLIM annotations (Gene Ontology Consortium et al, 2023) was calculated in both *S. cerevisiae* and *S. pombe*. For each set of orthologs, the number of ortholog pairs with more annotations in each species was calculated and the statistical significance was calculated by a binomial test. Then, the observed proportions of both sets were compared and the statistical significance was calculated by a Fisher’s exact test.

Next, ortholog gene pairs involving duplicated genes in either species were grouped by their interspecies interaction density deviation. First, the fraction of ortholog pairs involving duplicated genes in *S. pombe* with an interaction density deviation below -0.04 (lower *S. pombe* density than expected) was calculated. The same fraction was calculated for ortholog pairs involving duplicated genes in *S. cerevisiae*. Both fractions were compared and the statistical significance was calculated by a Fisher’s exact test. The same process was followed for deviation cutoffs of -0.03, -0.02, -0.01, and 0 to identify ortholog pairs with lower density in *S. pombe* than expected. Conversely, to identify cases with lower *S. cerevisiae* than expected, cutoffs of 0, 0.01, 0.02, 0.03, and 0.04 were used.

For N:M relationships, involving multiple genes in both species, the ortholog pairs were separated in two groups: i) cases in which there were more orthologs in *S. pombe*, and ii) cases in which there were more orthologs in *S. cerevisiae*. Next, the interaction density deviation was compared between both groups and the statistical significance was calculated by a Mann-Whitney U test.

### Comparison of paralog genetic interaction densities (1F, 1G)

Ortholog pairs with available genetic interaction data corresponding to orthology relationships with one gene in *S. pombe* and two genes in *S. cerevisiae* were selected. Pairs in which the *S. pombe* gene had very low negative genetic interaction density (lower than 0.01) were removed. Next, *S. cerevisiae* paralog pairs were grouped by their negative genetic interaction density in two sets: i) pairs in which both genes had a similar negative genetic interaction density in *S. cerevisiae*, being lower than expected given the *S. pombe* negative genetic interaction density and also lower than 0.05, ii) the rest of paralogs which include those with divergent and/or high negative genetic interaction density. For each paralog pair, the Jaccard similarity in cellular localization patterns (Huh et al, 2003) and their sequence identity was computed. Additionally, the sequence identity against the *S. pombe* single ortholog was also calculated. Then, these metrics were compared for both groups of paralogs and the statistical significance was evaluated by Mann-Whitney U tests.

For the second set of paralogs, including pairs with divergent and/or high negative genetic interaction density, the paralogs with the higher and lower negative genetic interaction density among each pair were compared using three different datasets: i) cellular localization similarity to the *S. pombe* single ortholog (Matsuyama et al, 2006); ii) expression similarity against the *S. pombe* single ortholog (Koch et al, 2012); and iii) number of *S. cerevisiae* strains (out of a panel of 1,011 (Peter et al, 2018)) with a loss-of-function mutation. Statistical significance was calculated by Mann Whitney U tests.

### Conservation of genetic interactions (2A, 2B, 2C)

Gene pairs in *S. cerevisiae* with genetic interaction data and with orthologs in *S. pombe* were selected. For gene pairs screened more than once in *S. cerevisiae*, the pair was considered to interact if 50% or more of the tested combinations resulted in a negative genetic interaction. *S. cerevisiae* gene pairs were defined as having a negative genetic interaction in *S. pombe* if any of the possible ortholog pairs in that species had a negative genetic interaction. For instance, if both *S. cerevisiae* genes had a single ortholog in *S. pombe*, then only one gene pair in that species was evaluated. Conversely, if both *S. cerevisiae* genes had multiple orthologs, then all possible ortholog pairs were evaluated. Next, *S. cerevisiae* gene pairs were split between those with and without a negative genetic interaction, and the fraction of cases with a negative genetic interaction in *S. pombe* was compared. Fold enrichment was calculated by dividing the ratio of *S. cerevisiae* negative genetic interactions with a negative genetic interaction in *S. pombe* by the corresponding ratio among *S. cerevisiae* pairs without a negative genetic interaction. Statistical significance was evaluated by a Fisher’s exact test. The same procedure was used for positive genetic interactions, and for gene pairs classified according to their orthology relationship in *S. pombe*: i) both genes have 1:1 orthologs; ii) only one of the genes have a 1:1 ortholog; iii) none of the genes have 1:1 orthologs.

### Conservation of genetic interactions by strength (2D)

Gene pairs were sorted by their interaction strength in *S. cerevisiae* and a precision recall curve was calculated for the identification of negative genetic interactions in *S. pombe*. Additionally, gene pairs with a negative genetic interaction in *S. cerevisiae* were selected, and the median epsilon score was used to define two sets of negative genetic interactions in that species: one with high genetic interaction scores (strong negative genetic interactions) and the other with low genetic interaction scores (weak negative genetic interactions). Then the fraction of corresponding negative genetic interactions in *S. pombe* for each set was compared. Statistical significance was evaluated by a Fisher’s exact test.

### Conservation of genetic interactions by functional relationship (2E)

Gene pairs in *S. cerevisiae* were annotated to five functional standards: coannotation if they had the same biological process GO annotation (Myers et al, 2006; Costanzo et al, 2016), colocalization if they were found in the same cellular compartment (Huh et al, 2003), coexpression if they displayed similar expression profiles (MEFIT score above 1.0) (Huttenhower et al, 2006), copathway if they were part of the same molecular pathway (Kanehisa et al, 2023), and cocomplex if they coded for proteins in the same protein complex (Meldal et al, 2021). Next, gene pairs annotated to each of the functional standards were selected, and the negative genetic interaction conservation across species was evaluated as explained above (see “Conservation of genetic interactions”). Finally, gene pairs were split by their number of functional annotations. Each set was then evaluated for negative genetic interaction conservation across species.

### Conservation of genetic interactions by broad functional classes (2F)

Gene pairs in *S. cerevisiae* were annotated to broad functional classes previously defined (Costanzo et al, 2016). Then, for each functional class, gene pairs in which both genes were annotated to that function were selected and evaluated for genetic interaction conservation as explained above (see “Conservation of genetic interactions”).

### Prediction of conserved genetic interactions across species (2G)

Gene pairs with a negative genetic interaction in *S. cerevisiae* and their functional relationships, genetic interaction scores, and orthology relationships in *S. pombe* were selected as explained above. This data was used to train a random forest (Liaw & Wiener, 2002) for the prediction of conserved negative genetic interactions in *S. pombe*. The resulting out-of-bag predictions (made with data not used for training) were sorted by their prediction score and a precision recall curve was calculated for the identification of conserved negative genetic interactions in *S. pombe*. The ratio of conserved negative genetic interactions for predictions within the top 1% recall was compared to the rest of predictions. Statistical significance was evaluated by a Fisher’s exact test. The same approach was followed for predictions within the top 10% and top 50% recall.

### Conservation of genetic interactions by ortholog similarity (2H)

Gene pairs with genetic interaction data in *S. cerevisiae* and 1:1 orthologs in *S. pombe* were selected. For each gene pair, the similarity with their orthologs was calculated using three different metrics: localization, protein-protein interactions and sequence. Similarity in localization was defined as both genes being localized in at least one same compartment as their orthologs (Huh et al, 2003; Matsuyama et al, 2006). Similarity in protein-protein interactions was defined as both genes and their corresponding orthologs having at least one common PPI. PPI data was downloaded from HINT (Das & Yu, 2012), and only PPIs against proteins with 1:1 orthologs were considered. Similarity in sequence was defined for the top 10% average sequence similarities between both genes and the corresponding orthologs. Then, for each gene pair the number of metrics in which they were similar to their orthologs was calculated. Gene pairs were split by the number of similar metrics and the negative genetic interaction conservation in *S. pombe* was calculated as explained above.

### Fitness correlation across species (3A)

Single mutant fitness (SMF) in *S. pombe* (Koch et al, 2012) was correlated against three different fitness sets in *S. cerevisiae* (Costanzo et al, 2016): i) SMF of 1:1 orthologs, ii) SMF of *S. cerevisiae* genes corresponding to orthology relationships with one gene in *S. pombe* and two in *S. cerevisiae*, and iii) the corresponding double mutant fitness in *S. cerevisiae* of the previous set. Only non-essential genes were considered in both species. Correlation was calculated using the Spearman’s correlation coefficient.

### Conservation of genetic interactions for gene duplicates (3C, 3D, 3E)

Gene pairs in *S. cerevisiae* in which one gene had a 1:1 ortholog and the other had a N:1 ortholog in *S. pombe* (i.e. duplicated in *S. cerevisiae*) were selected. The number of paralog genes in *S. cerevisiae* corresponding to the N:1 ortholog relationship was annotated. Pairs without a negative genetic interaction in *S. pombe* were disregarded. Next, *S. cerevisiae* gene pairs were grouped by the number of paralogs of the duplicated gene. The fraction of negative genetic interactions in *S. cerevisiae* was compared for the sets of *S. cerevisiae* gene pairs with i:) two, ii) three, and iii) four or more paralogs. Statistical significance was calculated by Fisher’s exact tests.

Following, the set of gene pairs with only two paralogs were selected such that each *S. pombe* gene mapped to two different *S. cerevisiae* gene pairs. Next, the sequence identity between the paralog pairs was calculated. Gene pairs were split by whether any of the two related gene pairs in *S. cerevisiae* had a conserved negative genetic interaction. Then, sequence identity was compared across the two sets and the statistical significance was evaluated by Mann-Whitney U tests. Finally, gene pairs in which one of the paralogs had a conserved negative genetic interaction were selected. Cases in which the conserved negative genetic interaction corresponded to the paralog most similar and most dissimilar in expression level to the *S. pombe* ortholog gene were counted. Statistical significance was evaluated by a binomial test.

### Negative trigenic interactions analysis (3F)

Gene pairs with genetic interaction data in *S. pombe* were selected in which one gene had a 1:1 ortholog and the other had a 1:2 ortholog relationship (i.e. two duplicates) in *S. cerevisiae*. Next, data from previous large-scale studies (Kuzmin et al, 2018, 2020) was used to identify negative trigenic interactions between the sets of three *S. cerevisiae* genes. The fraction of negative trigenic interactions was compared for the set of pairs with and without a negative genetic interaction in *S. pombe*. Statistical significance was evaluated by a Fisher’s exact test. The same approach was used for the subset of gene pairs in which the *S. cerevisiae* duplicates had a negative genetic interaction, and for the subset without a negative genetic interaction. The predictions of negative trigenic interactions were generated using gene pairs in *S. pombe* with a negative genetic interaction, in which the duplicated genes in *S. cerevisiae* had also a negative genetic interaction, and that were not screened for trigenic interactions in *S. cerevisiae*. **Table S2** includes 176 negative trigenic interaction predictions.

### Correlation of genetic interaction profile similarities (4A, 4B, 4C, 4E)

Genetic interaction profile similarities for *S. cerevisiae* were downloaded from TheCellMap (https://thecellmap.org/) (Usaj et al, 2017). In genes with data for more than one allele, the similarities were averaged across alleles. Genetic interaction profile similarities for *S. pombe* were computed using the Pearson’s correlation coefficient on the array profiles. A table was generated including all *S. cerevisiae* gene pairs with genetic interaction profile similarities and the corresponding ortholog pairs in *S. pombe*. Each possible ortholog pair was assigned a different row in that table. For instance, if both genes in *S. cerevisiae* had 1:1 orthologs in *S. pombe*, only one row would be generated. However, if both *S. cerevisiae* genes had two orthologs in *S. pombe*, then four rows would be generated. That table was used to calculate the general correlation between *S. cerevisiae* and *S. pombe* genetic interaction profile similarities. The Pearson’s correlation coefficient was calculated between the genetic interaction profile similarities of both species for: i) all gene pairs; ii) gene pairs in which both genes had a 1:1 ortholog; iii) colocalized gene pairs in *S. cerevisiae* (Huh et al, 2003); iv) gene pairs annotated to the same broad function in *S. cerevisiae* (Costanzo et al, 2016); v) gene pairs coding for proteins in the same molecular pathway (Kanehisa et al, 2023) or protein complex (Meldal et al, 2021) in *S. cerevisiae*.

The agreement among gene pairs with high similarity in both species was calculated by using the top 0.1% value in each similarity network. Then, gene pairs in *S. cerevisiae* were split by whether their similarity value was above that cutoff. For each set, the fraction of gene pairs in *S. pombe* with a similarity above the top 0.1% cutoff was calculated. Fold enrichment was calculated by dividing both fractions, and the statistical significance was computed by a Fisher’s exact test. The same approach was followed for the top 1% and top 10% similarity values.

Next, rows in the similarity table corresponding to gene pairs with 1:1 orthologs were selected. For each *S. cerevisiae* gene_x_ in the table, the gene_y_ with the most similar genetic interaction profile similarity was identified. Then, for the corresponding *S. pombe* ortholog of gene_x_, the genetic interaction profile similarities in *S. pombe* were sorted. The following cases were considered regarding gene_y_ ortholog and the sorted similarities in *S. pombe*: i) it was also the top 1 similarity; ii) it was among the top 1% similarities; iii) it was among the top 10% similarities. To calculate the background expectation, the same calculations were repeated using *S. cerevisiae* genes different than gene_y_. Fold enrichments were calculated by comparing the results of gene_y_ (the top 1 similarity in *S. cerevisiae*) to the background expectation. Statistical significance was calculated by Fisher’s exact tests.

### Correlation of genetic interaction profile similarities (4D)

Gene pairs with a genetic interaction profile similarity in *S. pombe* among the top 1% values were selected. The subset in which the *S. cerevisiae* profile similarity was also among the top 1% values in that species was defined as the set with conserved similarities across species. Conversely, the remaining gene pairs were defined as the set in which the high similarity in *S. pombe* was lost in *S. cerevisiae*. Then, for the conserved and lost sets, the fraction of gene pairs in which: i) both genes had 1:1 orthologs; ii) one gene had a 1:1 ortholog and the other was duplicated in *S. cerevisiae*; iii) both genes were duplicated in *S. cerevisiae* was calculated. Fold enrichment was calculated by dividing the ratios in the conserved set by the corresponding ratios in the lost set. Statistical significance was evaluated by Fisher’s exact tests.

### Interspecies genetic interaction profile correlation (5A)

For *S. cerevisiae*, array profiles corresponding to genes with 1:1 orthologs in *S. pombe* were selected. Only a query and array allele per gene was randomly selected. For *S. pombe*, array profiles corresponding to genes with 1:1 orthologs in *S. cerevisiae* were selected. Additionally, only array and query genes present in both networks were considered. This filtering process resulted in 484 array profiles including 267 query genes for each species. Next, interspecies array genetic interaction profile similarities were computed using the Pearson’s correlation coefficient between both sets of profiles. Similarities between 1:1 ortholog pairs and the rest of pairs were compared. Statistical significance was evaluated using a Mann-Whitney U test.

Next, the fraction of ortholog pairs with an interspecies genetic interaction profile similarity above 0.1 was compared to the corresponding fraction for non-ortholog pairs. Statistical significance was evaluated by Fisher’s exact test. The same process was performed for similarity cutoffs above 0.2 and 0.3.

### Characterization of orthologs with high profile similarity (5B, 5C)

Interspecies genetic interaction profile similarities for ortholog pairs were selected. Ortholog pairs within the top 10% similarities were compared against the rest of ortholog pairs using Qarles (Pons, 2025), a web server which evaluates 31 numeric and binary features to characterize gene sets in *S. cerevisiae*. Statistical significance was evaluated by Mann Whitney U tests and Fisher’s exact tests for numeric and binary features, respectively. Features significantly enriched with an FDR < 0.05 were reported. Additionally, the similarity in cellular localization between species (Huh et al, 2003; Matsuyama et al, 2006) was calculated using the Jaccard index, and compared for both sets of orthologs. Statistical significance was calculated by a Mann-Whitney U test. Finally, 15 broad functions in *S. cerevisiae* (Costanzo et al, 2016) were used to annotate both sets of orthologs. For each function, the fold enrichment was calculated by dividing the ratio of similar orthologs annotated to that function by the corresponding ratio for the rest of orthologs. Statistical significance was calculated by Fisher’s exact tests.

### Characterization of non-ortholog pairs with high profile similarity (5D)

Interspecies genetic interaction profiles similarities between non-ortholog pairs were selected. Gene pairs with similarity above 0.3 were compared to the rest of pairs by using cocomplex membership annotations in *S. cerevisiae*. The fold enrichment was calculated by dividing the fraction of similar gene pairs annotated to cocomplex associations (Meldal et al, 2021) by the corresponding ratio for the rest of gene pairs. Statistical significance was calculated by Fisher’s exact tests. The same approach was used for similarity cutoffs above 0.2, and 0.1, and with copathway (Kanehisa et al, 2023), colocalization (Huh et al, 2003), and same broad function annotations (Costanzo et al, 2016).

### Interspecies genetic interaction profile similarity of protein complexes (5E)

Protein complexes in *S. cerevisiae* with 2 or more subunits (Meldal et al, 2021) with available interspecies similarities were selected. For each complex, the interspecies similarities between all possible gene pairs were retrieved, including the ortholog pair. Statistical significance was evaluated by Mann-Whitney U tests against all interspecies similarities and corrected for multiple tests by using the FDR.

